# The motor neuron-like cell line NSC-34 and its parent cell line N18TG2 have glycogen that is degraded under cellular stress

**DOI:** 10.1101/2020.03.23.004184

**Authors:** Brigitte Pfeiffer-Guglielmi, Ralf-Peter Jansen

**Affiliations:** Interfaculty Institute for Biochemistry, University of Tübingen, Hoppe-Seyler-Str. 4, D-72076 Tübingen, Germany

**Keywords:** Neuronal cell lines, glycogen, glycogen phosphorylase isozymes, hypoxia stress, oxidative stress, pH-stress

## Abstract

Brain glycogen has a long and versatile history: Primarily regarded as an evolutionary remnant, it was then thought of as an unspecific emergency fuel store. A dynamic role for glycogen in normal brain function has been proposed later but exclusively attributed to astrocytes, its main storage site. Neuronal glycogen had long been neglected, but came into focus when sensitive technical methods allowed quantification of glycogen at low concentration range and the detection of glycogen metabolizing enzymes in cells and cell lysates. Recently, an active role of neuronal glycogen and even its contribution to neuronal survival could be demonstrated. Our studies continue these investigations on the function and regulation of neuronal glycogen metabolism. We demonstrate the presence of an active glycogen metabolism in the neuronal cell lines NSC-34 and N18TG2 and the mobilization of the glycogen stores under hypoxia, oxidative and acidic metabolic stress. The key enzyme in glycogen degradation is glycogen phosphorylase. Neurons express only the brain isoform (GPBB) that is supposed to be activated primarily by the allosteric activator AMP and less by covalent phosphorylation via the cAMP cascade. Our results indicate that neuronal glycogen is not degraded upon hormone action but by factors lowering the energy charge of the cells directly.

## Introduction

Glycogen represents the major brain energy reserve though its quantities are small compared to muscle and liver. Because neural glycogen is predominantly located in astrocytes, its possible functions have been attributed for decades to these cells with the consensus that glycogen is an emergency fuel reserve for conditions of physiological or pathological stress. However, more recent studies have shown that glycogen serves a dynamic role also in normal brain metabolism, e.g. in neurotransmission, learning and memory consolidation, and in sleep (1,2,3,4).

A physiological role of glycogen in CNS neurons has long been denied though they express glycogen synthase and have the potential to synthesize glycogen under pathological conditions (5). In neurons of the PNS and in spinal motor neurons, the intense immunoreactvity for the brain isoform of glycogen phosphorylase (GPBB) made the presence of its substrate highly probable (6,7). The current view on neuronal glycogen changed with the availability of highly sensitive methods allowing the quantitative determination of glycogen in cortical neuronal cultures (8) as well as the detection of the glycogen metabolizing enzymes at the protein and mRNA levels (9). Though the glycogen content of cortical neurons only mounts up to about 1/10 of that of astrocytes, it could be demonstrated that neurons have an active glycogen metabolism that contributes to tolerance to hypoxia (8). These findings have drawn new attention to the role of neuronal glycogen in health and disease (10,11,12).

Under these aspects, the high immunoreactivity for GPBB in peripheral neurons and spinal motor neurons may be an indicator for an active glycogen metabolism in these cells. The high vulnerability of motor neurons and their involvement in neurodegenerative diseases like amyotrophic lateral sclerosis (ALS) make these cells a highly probable candidate for glycogen utilization. Metabolic studies like glycogen determination under different metabolic conditions, however, require cell culture systems with high yields and purity. Peripheral neurons like DRG neurons as well as spinal motor neurons can be cultured, the latter even in high purity, but the cell yields are too low for metabolic studies. Neuronal cell lines, in contrast, would fulfill both conditions. NSC-34 is a hybrid cell line generated by the fusion of mouse embryonic motor neuron-enriched spinal cord cells with mouse neuroblastoma N18TG2 (13). This cell line is a relevant model for the study of motor neuron biology and expresses many motor neuron-like properties (for overview, see 13, 14,15). If NSC-34 had glycogen and the glycogen metabolizing machinery, these properties could originate from the primary parent cell (spinal cord motor neurons) or the parent tumor cell line (neuroblastoma). If NSC-34, but not N18TG2 had glycogen metabolizing properties, NSC-34 would be an ideal model for investigating motor neuronal glycogen. If both cell lines had an active glycogen metabolism, a general function for glycogen in neurons would become more likely. On the other hand, possible differences in glycogen content and/or glycogen mobilization under conditions of metabolic stress could indicate specific roles for glycogen in motor neurons and neuronal tumor cells.

In this study, we first investigated whether NSC-34 and N18TG2 contain glycogen and express the key enzymes of glycogen metabolism, glycogen synthase (GS) and glycogen phosphorylase (GP). Next, we looked for factors influencing glycogen metabolism in order to get insight into the metabolic regulation of glycogen degradation. The contribution of neuronal glycogen to tolerance of hypoxia stress makes glycogen a possible player in conditions of metabolic stress in general. We therefore examined glycogen mobilization in NSC-34 and N18TG2 cell lines under conditions of hypoxia, oxidative stress and pH lowering.

The enzyme catalyzing the rate-limiting step in glycogen degradation is glycogen phosphorylase. GP exists in three isoforms, GPBB, GPMM, GPLL, named according to the tissues they dominate in : brain, skeletal muscle, and liver. Astrocytes express GPBB and GPMM in equal amounts and perfect colocalization (6). Neurons express only the brain isoform (6,7). Consequently, glycogen degradation in neurons must be attributed to GPBB and metabolic effects on glycogen content found in neurons should also be found in astrocytes because of their GPBB fraction. We therefore included astrocyte primary cultures in our stress experiments. In addition, we investigated whether siRNA knockdown of GPBB blocks a possible stress-induced glycogen breakdown.

## Results

### NSC-34 and N18TG2 cell lines express GP and GS, the key-enzymes of glycogen metabolism and contain glycogen

In order to investigate whether the neuronal cell lines NSC-34 and N18TG2 have the enzymatic machinery to synthesize and degrade glycogen, we performed Western blotting. Both cell lines express GP and GS (Figure 1. A,B). Differentiation of NSC-34 does not influence the expression patterns of the enzymes (Figure 1.C). Both cell lines do not express GP MM (data not shown).

**Figure 1.**
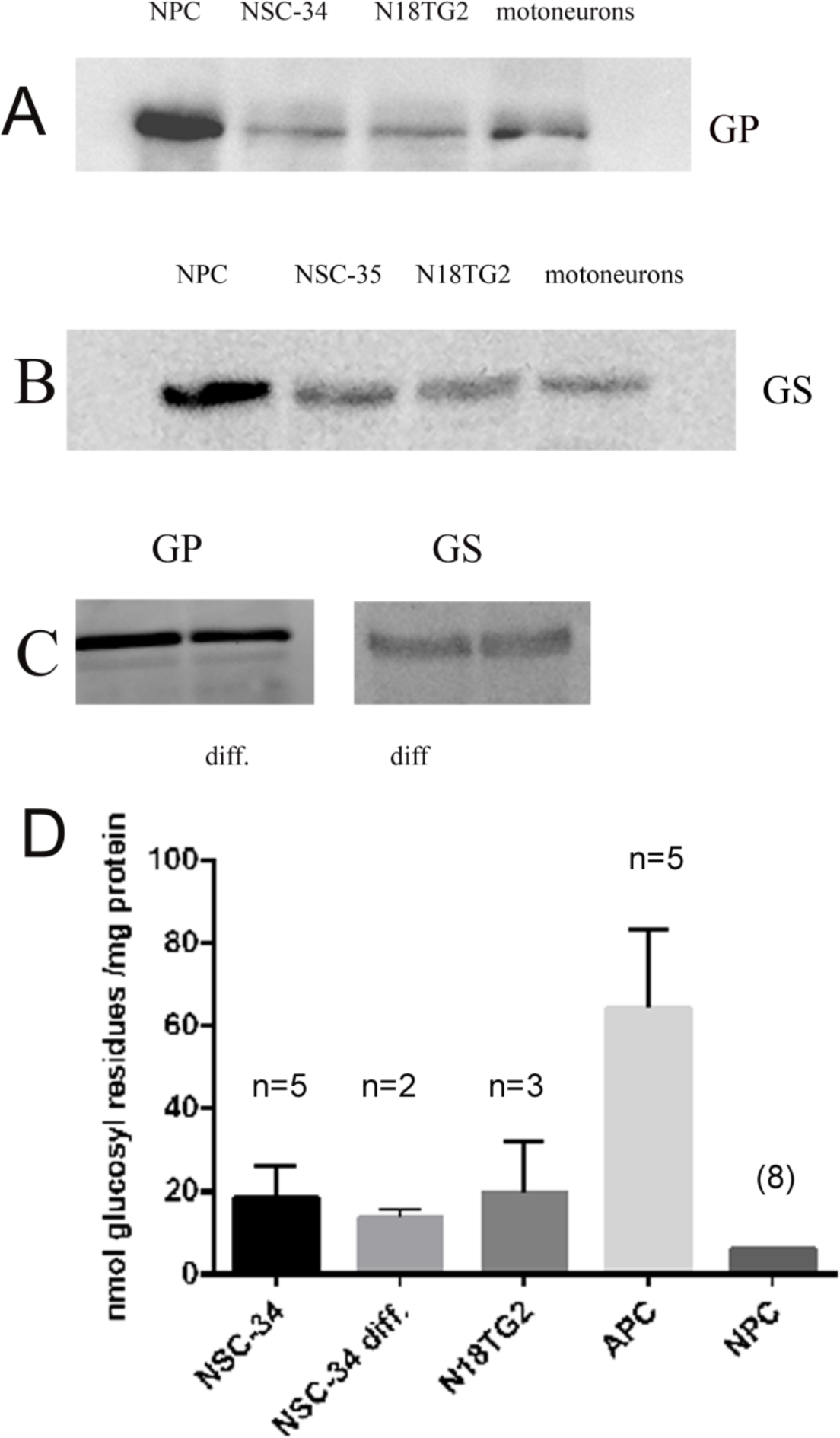
Western blot analysis for the detection of GPBB (A) and glycogen synthase (GS, B) in homogenates of NSC-34 and N18TG2 cells compared to NPC and pure motoneuron cultures. (C) GPBB and GS in differentiated NSC-34 cultures. Equal amounts of protein were applied in each blot. (D) Quantitative determination of glycogen in mouse neural cell types under the relevant culture conditions. Data represent mean values ± SD of n experiments each performed in triplicates.

We determined the glycogen content of confluent NSC-34 and N18TG2 cells under culture conditions (DMEM, 10 % FCS, 25 mM glucose). Glycogen values varied widely between experiments, but were comparably high in both cell types (Figure 1. D). They exceeded those found for primary neuronal cultures. Differentiation of NSC-34 (3 % FCS, 25 mM glucose) did not influence the glycogen content.

### The glycogen content of NSC-34 cells is dependent on the glucose concentration, but not on the presence of FCS

Because 25 mM glucose and the presence of 10 % FCS (culture conditions) do not reflect metabolic conditions, we examined the influence of glucose concentration and FCS on the glycogen content of NSC-34 neurons. In contrast to primary neuronal cultures (8), we found a decline in glycogen content with declining glucose concentration (Figure 2. A). Absence of FCS did not significantly alter the glycogen content (Figure 2. B) as also demonstrated in differentiated NSC-34 cultures (data not shown).

**Figure 2.**
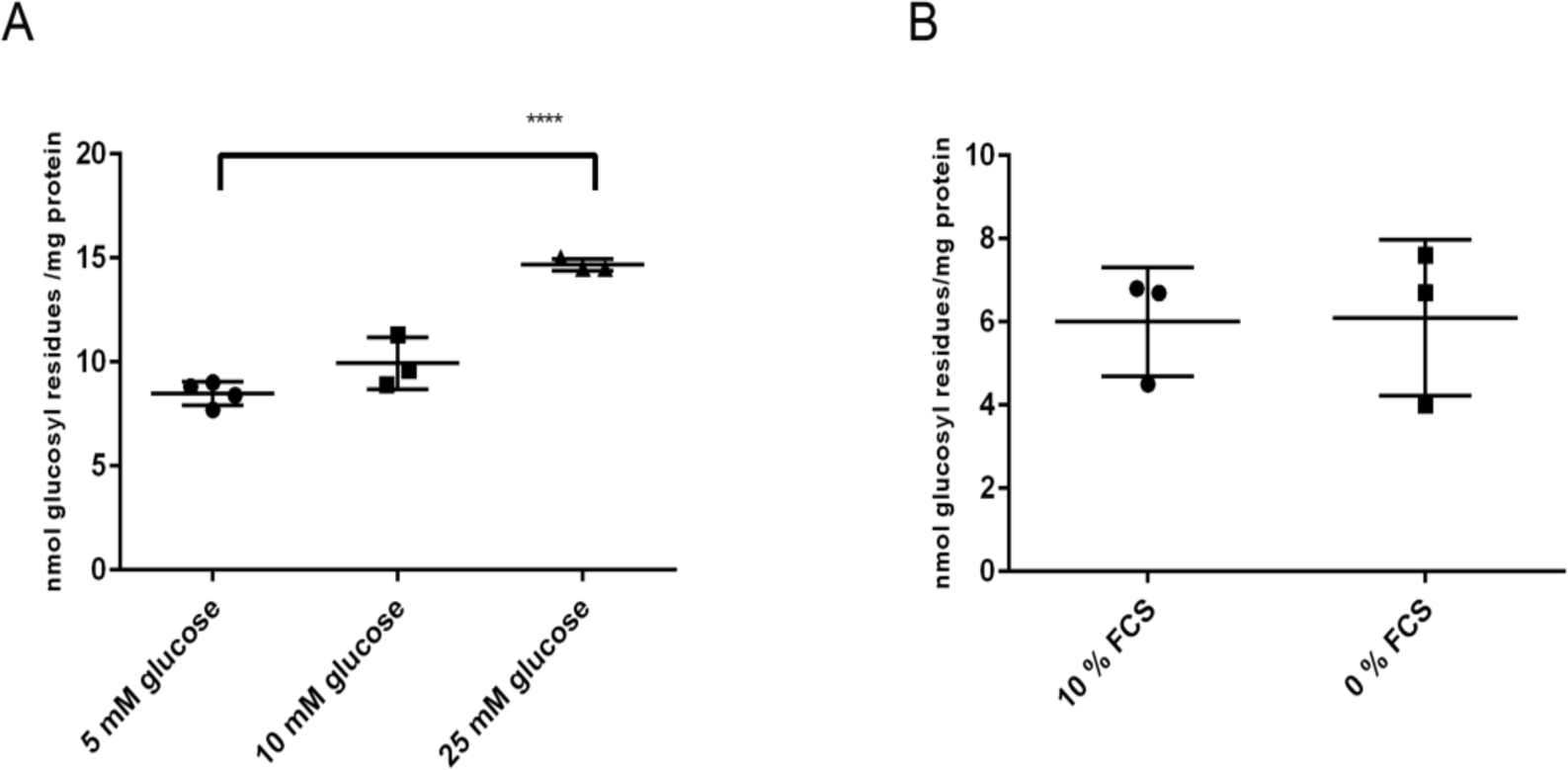
Influence of glucose concentration (A) and FCS (B) on the glycogen content of NSC-34 cultures. (A) Cells were cultured to confluency in DMEM/10 % FCS/ 25 mM glucose, then medium was changed to media containing 5, 10 and 25 mM glucose, respectively, and cells cultured for another 60 min. (B) Cells were cultured to confluency, then switched to DMEM/25 mM glucose with or without FCS and cultured for another 60 min. Significant differences are as indicated: **** p< 0,0001.

### Glycogen in NSC-34 cells is readily mobilizable by glucose deprivation, but not by hormones stimulating cAMP formation and by methoxamine

Next, we studied the dynamics of NSC-34 glycogen stores under glucose deprivation, a metabolic situation in which the AMP concentration raises. After 2 h on glucose-free DMEM, the glycogen store is depleted but can be replenished by a subsequent 2 h incubation with 5 mM glucose. This repletion results in a glycogen content exceeding the basal value (Figure 3. A).

**Figure 3.**
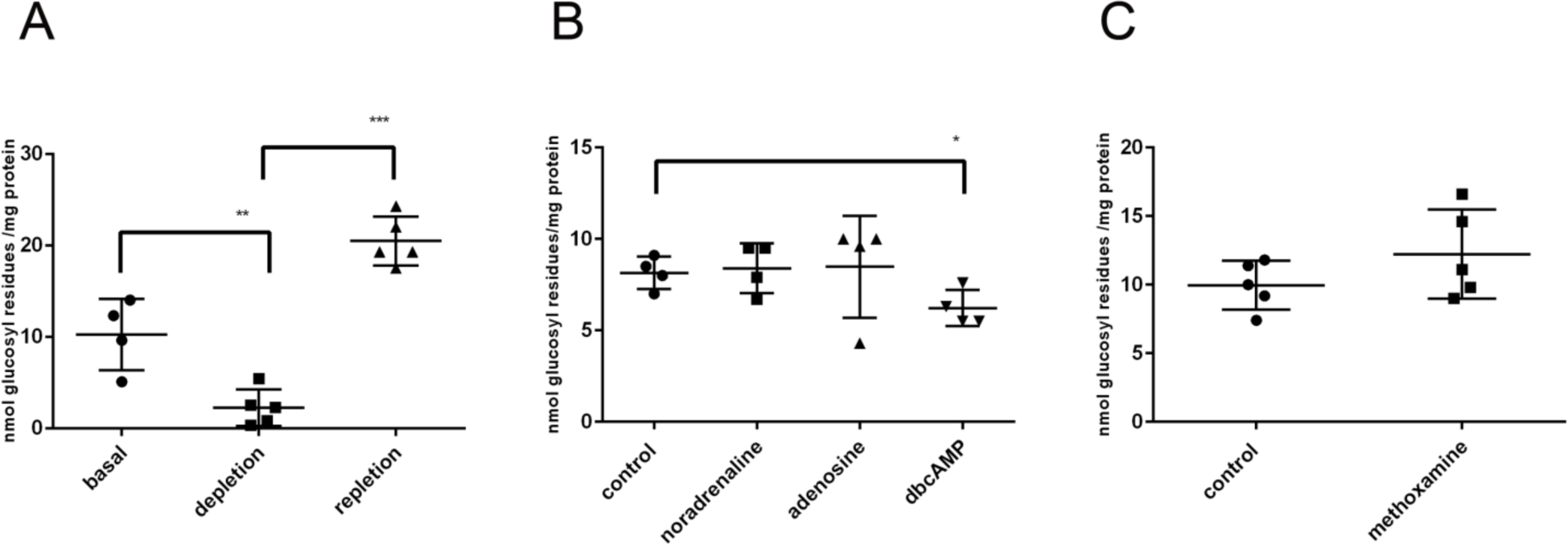
Influence of glucose deprivation (A) and modulators (B,C) on the glycogen content of NSC-34 cells. (A) Confluent cells were switched to glucose-free DMEM/ – FCS and cultured for 12 h (depletion). For repletion, cells were switched to DMEM/25 mM glucose/ -FCS for 2 h. (B,C) Confluent cells were switched to DMEM/5 mM glucose/ -FCS for 4 h. Then, modulators were added to a concentration of 100 μM and cells cultured for 30 min. Significant differences are as indicated: *p<0,05; **p<0,01; ***p<0,001.

In cultured astrocytes, glycogen is mobilized by vasointestinal peptide, noradrenaline, adenosine and dibutyryl-cAMP (16). Noradrenaline and adenosine did not lower the glycogen content of NSC-34 cells. Dibutyryl-cAMP, a membrane-permeable stable cAMP analog, elicited a moderate decline in glycogen content (Figure 3. B). Methoxamine, an activator of the phosphatidyl-inositol cascade, has also been shown to promote glycogenolysis in cultured astrocytes (17). In NSC-34 cells, methoxamine does not significantly lower the glycogen content (Figure 3. C).

### The glycogen content of NSC-34 neurons is influenced by the K^+^ -concentration

Neuronal activity and some pathological situations stimulate glycogenolysis in brain slices by raising extracellular [K^+^] and intracellular [Ca^2+^] (18,19). This glycogenolysis had been attributed to astroglial cells. In order to look whether a high K^+^ concentration influences neuronal glycogenolysis in cultures, we exposed NSC-34 cells to K^+^ concentrations reached at neuronal activity (10 mM) or under pathological conditions (40 mM). The glycogen content was lowered slightly at [K^+^] of 10 mM and significantly at 40 mM (Figure 4.).

**Figure 4.**
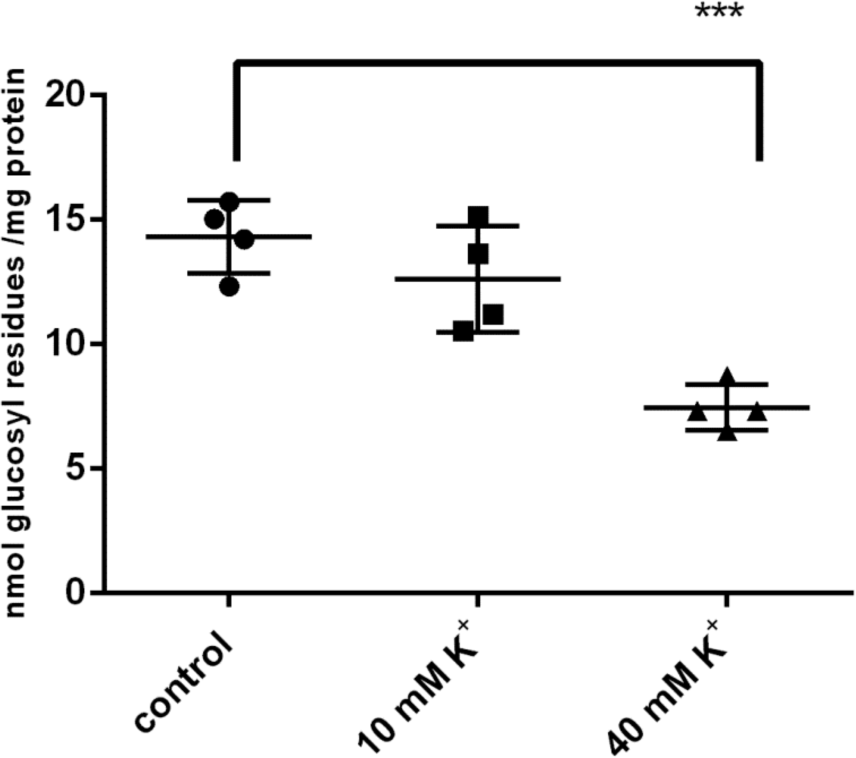
Infuence of elevated [K^+^] on the glycogen content of NSC-34 cells. Confluent cells were cultured in DMEM/ 5 mM glucose/ -FCS for 4 h and then switched to the indicated [K^+^] for 30 min. Significant differences are as indicated: ***p<0,001.

### Cellular metabolic stress induces glycogen degradation in NSC-34 and N18TG2 neurons

Under hypoxic conditions (1 % O_2_ instead of 21 %), cultured primary neurons degrade their glycogen stores by the action of glycogen phosphorylase (8). In order to investigate whether this might also apply to NSC-34 and N18TG2 cells, we exposed the cultures to hypoxic conditions. After 4 h of hypoxia, both cell lines showed a significant reduction of their glycogen content compared to normoxic conditions (Figure 5. A,B). The viability of the cells was 100 %.

In astroglia-rich primary cultures, glycogen is degraded following administration of peroxides (20). In order to look whether this metabolic stress situation is also a trigger for neuronal glycogen degradation, we exposed the cultures to mild oxidative stress (500 μM H_2_O_2_ for 30 min and 2 h respectively.). In NSC-34 and N18TG2, glycogen was reduced in a time-dependant manner (Figure 5. C,D). Viability was not influenced.

**Figure 5.**
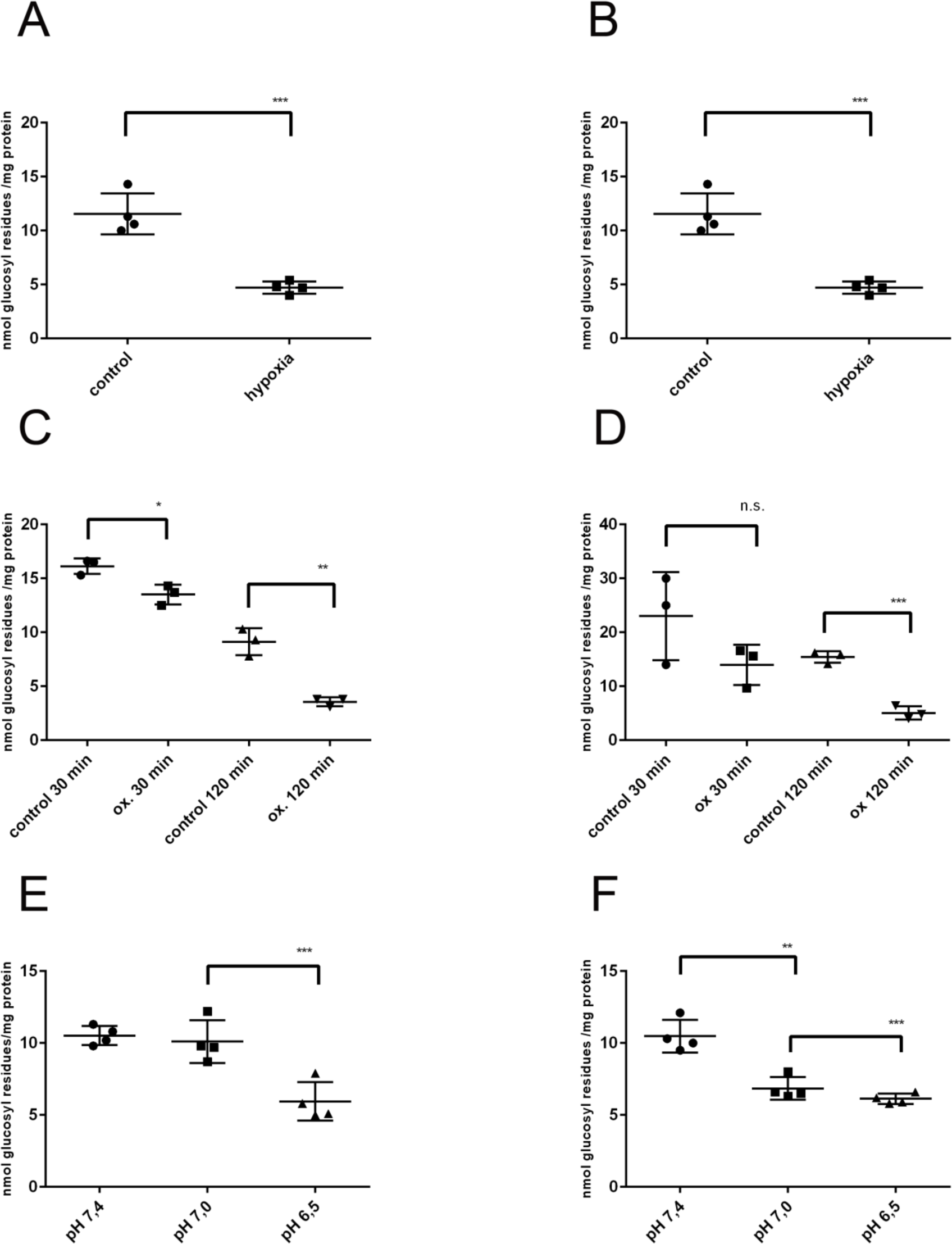
Influence of metabolic stress on the glycogen content of NSC-34 (A,C,E) and N18TG2 (B,D,F) cells. Confluent cells were switched to DMEM/5 mM glucose/ -FCS for 4 h and then exposed to the stress situations. (A,B) Hypoxia stress: Cells were exposed for 4 h to hypoxic conditions (1 % O_2_, 5 % CO_2_, 94 % N_2_), Control cells were kept under normoxic conditions (21 % O_2_). (C,D) Oxidative stress: H_2_O_2_ was added to a concentration of 500 μM and cells kept for 30 and 120 min, respectively. (E,F) Acidic pH: Cells were incubated for 1 h on a 37 °C water bath in a buffer containing 20 mM HEPES with Earle’s salts and 5 mM glucose, adjusted to the pH values indicated. Significant differences were as indicated: **p<0,01; ***p<0,001.

Deviation from the physiological value of pH is another form of metabolic stress. We therefore exposed the cell lines to moderate acidic pH conditions. Lowering the pH from 7,4 to 7,0 did not result in glycogen degradation. Acidification to pH 6,5, however, reduced the glycogen content in NSC-34 and in N18TG2 (Figure 5. E,F). Viability was slightly reduced at pH 6,5 (95 %).

### Astrocyte primary cultures and metabolic stress

Astrocytes in culture react with glucose degradation on hypoxia and oxidative stress analogously to neuronal cell lines (Figure 6. A, B). Lowering the pH from 7,4 to 7,1 leads to glycogen degradation, but in contrast to the cell lines, further lowering the pH to 6,5 reconstituted the glycogen store (Figure 6. C).

**Figure 6.**
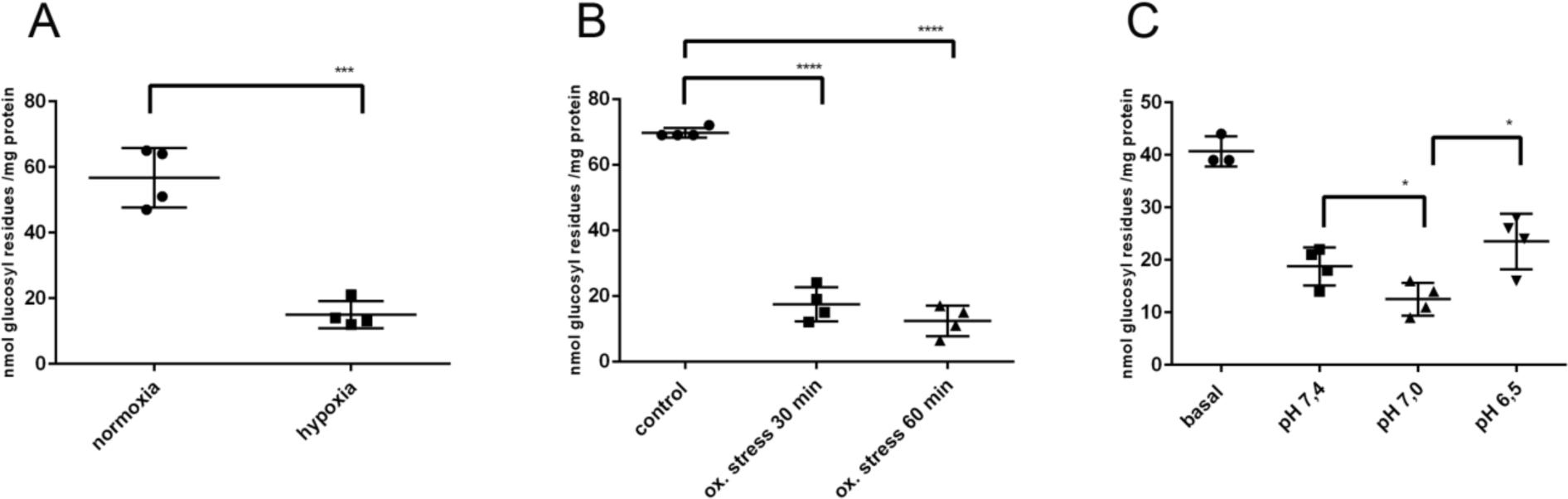
Influence of metabolic stress on the glycogen content of APC. (A) hypoxia; (B) oxidative stress; (C) Acidic pH. Cultures were used for experiments after two weeks of culturing and experiments carried out as described for the cell lines. Significant differences were as indicated: *p<0,05; **p<0,001; ***p<0,0001.

### Knockdown of GP BB abolishes glycogen degradation under metabolic stress

In order to demonstrate that the glycogenolytic answer to metabolic stress is due to GPBB action, we performed siRNA knockdown of GPBB in NSC-34 cells and APC with subsequent exposure to oxidative stress. After GPBB knockdown, glycogen content was elevated compared to untransfected cells (Figure 7. A,B). In cells transfected with targeting siRNA, glycogen was not reduced during exposure to oxidative stress. Cells transfected with NT siRNA, however, showed reduced glycogen content under oxidative stress (Figure 7 A,B). The Western blot signal for GPBB was reduced in cells transfected with siRNA (Figure 7. C,D).

**Figure 7.**
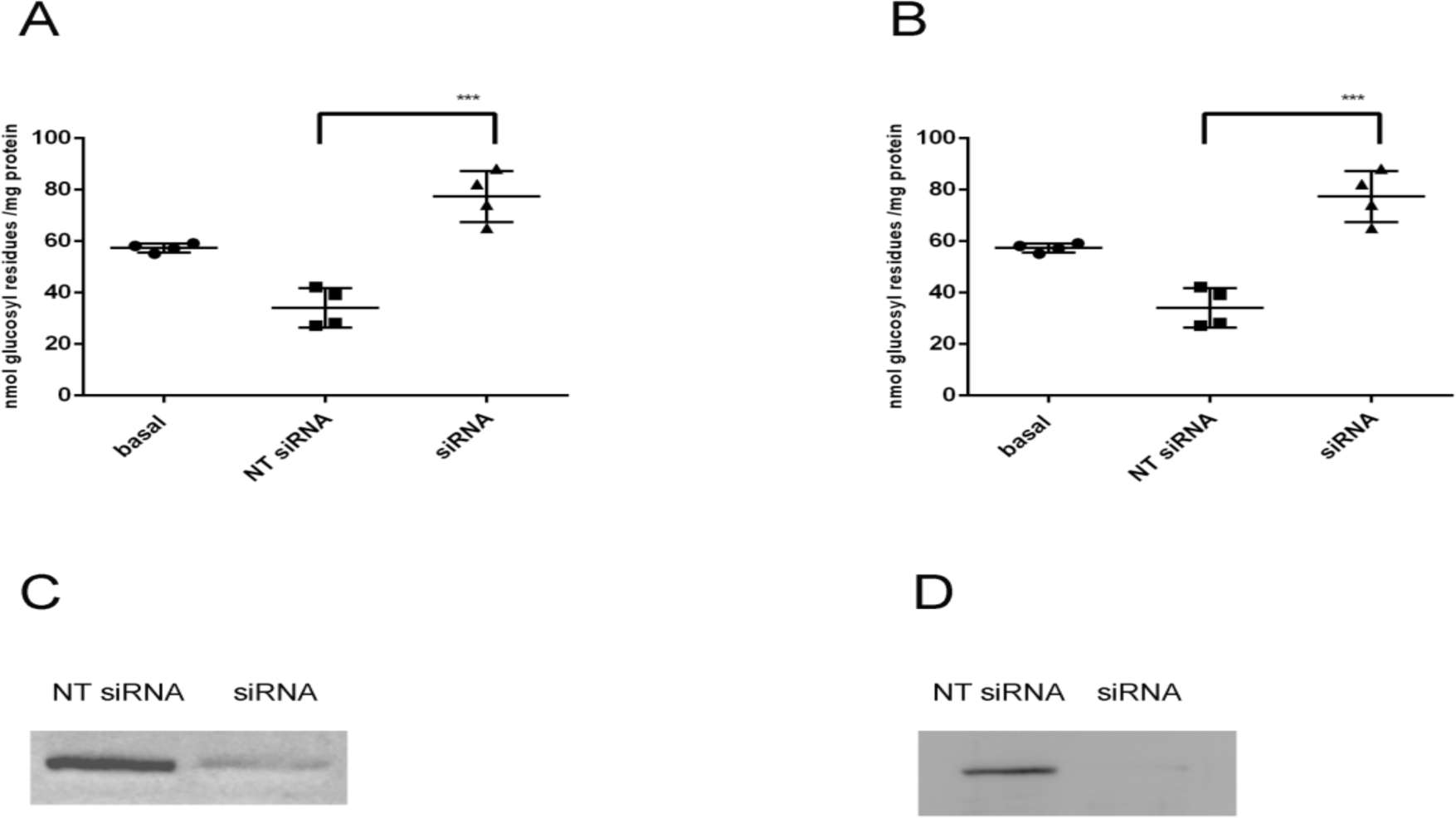
siRNA knockdown of GPBB with subsequent exposure to oxidative stress. (A) NSC-34 cells. (B) APC. (C,D) Western blot analysis of cell lysates of NSC-34 (C) and APC (D) at day 6 after transfection. Significant differences are as indicated: ***p<0,001.

## Discussion

After decades of neglecting a relevant role for neuronal glycogen besides that under pathological conditions, we could demonstrate the presence of glycogen and an active glycogen metabolism in the neuronal cell lines NSC-34 and N18TG2. In these cells, glycogen is degraded under conditions of metabolic stress thus confirming the results of Saez et al. (8) with cortical neuronal cultures.

Because of the high immunoreactivity of GP in spinal motoneurons (6,7,9) it seemed a logical idea that a cell line derived from these cells might have a higher glycogen content than cortical neurons. This was indeed the case: In the hybrid cell line NSC-34 we measured a glycogen content that was three-fold higher than that of cortical neurons. NSC-34 cultures comprise two morpholocically distinct populations: Neuroblastic cells with minimal projections and cells that undergo differentiation and have long processes (14). Differentiation can be achieved by cultivation under low serum conditions which is accompanied by high cell loss. Several authors use differentiated cells because some cell properties are only expressed in differentiated cells. In our experiments, differentiation did not influence neither the enzymatic pattern of GP and GS nor the glycogen content indicating that glycogen metabolism is not a privilege of differentiated cells.

The parent tumor cell line N18TG2 had a similar glycogen content and enzymatic machinery than the motoneuron/tumor cell hybrid, suggesting that glycogen metabolism is not an exclusive genetic contribution of the motoneuron parent. This pleads for a general importance of neuronal glycogen. Glycogen content of the neuronal cell lines exceed that of cortical neurons. This may be due to the specific metabolic situation of spinal motoneurons as well as of tumor cells.

In order to elucidate the function(s) of neuronal glycogen, we studied the influence of metabolic stress situations on the glycogen content. The simplest form of metabolic stress is glucose deprivation. We could demonstrate that the glycogen is rapidly mobilized under glucose deprivation, and that the store is replenished after glucose repletion showing a super compensation already demonstrated for cortical neurons after hypoxia (8).

Another situation of stress is hypoxia. It has been demonstrated that under hypoxic conditions, glycogenolysis occurs though extracellular glucose concentration is high, and prevents cortical neurons from cell death (8). Under these conditions, cellular uptake of glucose and its metabolization via glycolysis is obviously not sufficient to meet the requirements of the neurons. Degradation of glycogen, in contrast, leads to a faster energy availability. We found that NSC-34 and N18TG2 but also APC, reacted upon hypoxia with glycogen mobilization.

Reactive oxygen species (ROS) are continuously generated in cells during oxidative metabolism. Elevated levels of ROS are a hallmark in nerve cell injuries because they can lead to cell death via diverse oxidative and peroxidative reactions (21,22). It has already been demonstrated that in cultured astrocytes, glycogen was mobilized during the disposal of peroxides (20). This glycogenolysis was interpreted as a source for glucose-6-phosphate, the substrate fueling the pentose phosphate shunt which is necessary for the generation of NADPH. NADPH, in turn, is needed for the glutathione redox cycling system in cellular peroxide detoxification. Our findings that mild oxidative stress led to glycogenolysis in neuronal cell lines, might be a hint to a similar mechanism in highly sensitive neurons.

Another situation of metabolic stress is acidification. Mainteinance of the extracellular pH within strict boundaries is crucial for normal cell function. Extracellular acidity is observed in ischemia and acute inflammation, but is also a hallmark of solid tumors giving those cells an advantage over normal cells. It could be demonstrated that in HEK cells extracellular pH lowering induced mitochondrial dysfunction, protein carbonylation and increased ROS levels (23). In melanoma cells, extracellular acidity increases intracellular Ca^2+^ levels (24). We could find that in NSC-34 and N18TG2 cells pH lowering to 6.5 resulted in glycogenolysis. Neuronal excitability and neurotransmission induce pH alterations which, in turn, regulate pH sensitive transport systems (25) involving energy consumption. Rapid energy supply by glycogen degradation might play a role in situations of extracellular acidosis. Also, elevated ROS levels might request activation of the pentose phosphate shunt. In addition, elevated intracellular [Ca^2+^] can activate GP by a Ca^2+^ -dependant mechanism as demonstrated in brain slices (18). In APC, we observed glycogenolysis upon pH lowering from 7,4 to 7,0. Lowering the pH to 6,5, however, resulted in a reconstitution of the glycogen content to values exceeding that at pH 7,4, a super-compensation that may serve as an anticipatory mechanism of further stress.

We tried to get insight into the mechanism by which glycogen degradation is triggered in neurons. GP can be activated by two mechanisms, 1) covalent phosphorylation triggered by hormones via the cAMP cascade, and 2) allosteric control by AMP. The MM isoform is primarily activated by the first mechanism, while the BB isoform is more susceptible to the second mechanism (26). Selective knockdown of the two isoforms in cultured astrocytes confirmed these differences (27). We found that in neuronal cell lines neither noradrenaline nor adenosine lowered the glycogen level. dbcAMP, a cell-permeable cAMP analog that circumvents the receptor binding step and acts directly on protein kinase A, shows a moderate effect. IP_3_ can also activate glycogenolysis by binding to α_1_-receptors which leads to activation of proteinkinase C and elevated intracellular Ca^2+^-levels. In our cell models, methoxamine, an activator of the IP_3_-induced glycogenolysis in astrocytes (17), did not show a significant effect. Glucose deprivation, however, led to glycogenolysis. Knockdown of GPBB reduced the stress-induced glycogenolysis. These results indicate that neuronal glycogenolysis is primarily induced by the energy status of the cell.

Neuronal activity raises the extracellular K^+^ concentration to 5 – 12 mM. As could be demonstrated with cortical slices, this leads to the opening of voltage-gated Ca^2+^ - channels and elevated intracellular Ca^2+^ - concentrations resulting in glycogenolysis (18). Under pathological conditions like hypoxia, spreading depression or ischemia, extracellular K^+^ - concentrations up to 40 mM have been reported (19). In cultured astrocytes, glycogenolysis is triggered by store-operated Ca^2+^ entry (27). In NSC-34 cells, elevation of extracellular K^+^ resulted in glycogenolysis. probably by a Ca^2+^ - dependent mechanism. Ca^2+^ stimulates glycogenolysis by activation of phosphorylase kinase. Because GPBB is weakly activated by phosphorylation, elevated cytosolic Ca^2+^ might induce glycogenolysis by increasing the AMP concentration. This is the case when energy consumption and thus ATP hydrolysis take place. ATP is needed for maintaining ion gradients after Ca^2+^ influx (27). In this metabolic situation, glycogenolysis is the fastest way to replenish ATP stores.

Here, we could demonstrate the presence of glycogen and an active glycogen metabolism in neuronal cell lines. Because we found no difference between the motor neuron/ neuroblastoma hybrid NSC-34 and the tumor cell line N18TG2, the glycogen metabolizing machinery seems not to be a unique tool of spinal cord motor neurons. In our models, glycogenolysis is elicited by situations of metabolic stress thus pleading in favor of a so far neglected relevant role of neuronal glycogen in general. Neurons express, in contrast to astrocytes, only the GPBB isoform which is supposed to be primarily activated by the AMP level. In our studies, hormones activating covalent phosphorylation of GP, had no glycogenolytic effect. This supports the proposed functional differences of the GP isoforms.

## Experimental procedures

### Cell culture

The NSC-34 cell line was purchased from Cedarline (Burlington, Ontario, Canada) via tebu-bio (Offenbach, Germany) and cultured in DMEM/ 10 %FCS/ 25 % glucose with penicillin and streptomycin. For studies with differentiated cells, cultures were switched to a medium with low FCS concentration (DMEM/ Ham’s F 12 1:1, 1 % Eagle’s medium NEAA/ 3 % FCS/ 25 mM glucose with penicillin and streptomycin, 28). The N18TG2 cell line was purchased from the German Collection of Microorganisms and Cell Cultures (DSMZ, Braunschweig, Germany) and cultured in DMEM/10 % FCS/ 100 μM thioguanine / 25 mM glucose. Both cell lines were cultured in 60 mm plates at 10 % CO_2_ and 37 °C until confluency.

Neuronal primary cultures (NPC) and astrocyte primary cultures (APC) were prepared as described (9).

### siRNA knockdown

NSC-34 cells were cultured to 80 % confluency and transfected with ON-TARGET plus siRNA (Dharmacon) targeting the GPBB sequences as described (27). FUGENE HD Transfection Reagent (Promega, Madison, Wi, USA) was used according to the manufacturer’s protocol. siRNAs were applied in a concentration of 25 nM in OptiMEM medium (Gibco). APC were transfected analogously in the second week in culture. 24 h post transfection, medium was changed to culture medium and cells were cultured for 5 days. Negative controls were performed by applying a non-targeting siRNA (Dharmacon). Knockdown was controlled by Western blotting.

### Western Blotting

Supernatants of lysates from NSC-34, N18TG2, NPC and APC were prepared as described (9). Supernatants of lysates from motoneuron cultures were prepared in RIPA buffer (50 mM Tris/HCl pH 7,4; 150 mM NaCl, 1 % Triton X-100 and 2 mM EDTA) and were a kind gift of S. Jablonka (Würzburg). Western blotting was performed as described (9). In addition to the peroxidase/chemiluminescence detection system, we applied IRDye® 680 RD goat anti-rabbit antibody (LI-COR-Biosciences, Bad Homburg, Germany). Rabbit antisera against GP BB and GP MM were prepared as described (6). Rabbit monoclonal antibody against glycogen synthase (mAb 15B1) was purchased from Cell signaling Technology (Danvers, Mas., USA).

### Protein determination

Protein was determined using a microtiter plate test with Roti-Quant reagent (Roth, Karlsruhe, Germany) and bovine serum albumin as standard.

### Glycogen determination

Glycogen content of cell cultures was determined applying a method described (8) in combination with a fluorometric assay (29). After incubation of the cells under the relevant metabolic conditions, the culture medium was removed and the cells rapidly frozen and stored at −80 °C. For analysis, cells were scraped off the plates in a volume of 400 μl of 30 % KOH and the lysate incubated for 15 min at 100 °C. After taking an aliquot for protein determination, glycogen was precipitated in the lysate by the addition of ethanol to a concentration of 65 % (v/v) overnight at −20 °C. After centrifugation at 12 000 g for 10 min at 4 °C, pellets were air-dried overnight at room temperature. Thereafter, glycogen was digested in a volume of 200 μl of 50 mM acetate buffer pH 4,8 with 1 U of amyloglucosidase (1h, 37 °C). Then, glucose was determined with a fluorometric microplate assay under the following conditions: 20 μl of the amyloglucosidase digest were incubated for 30 min at 37 °C with 80 μl of 100 μM triethanolamine buffer pH 7,6 containing 1 mM MgCl_2_, 1 mM ATP, 100 μM NADP, 10 μM resazurin, 0,1 U/ml glucose-6-phosphate dehydrogenase, 0,2 U/ml diaphorase and 12 U/ml hexokinase. Fluorescence intensity was measured at an excitiation wavelength of 530 nm and an emission wavelength of 590 nm. Glycogen was expressed as nmol glucose residues and normalized to the protein content of the culture plates.

### Metabolic studies

For studying the influence of various parameters on the glycogen content, the cells were cultured to confluency and then switched to the conditions of the relevant experiment (for details, see Results).

To study the glycogen content under conditions of cellular stress, confluent cultures were switched to DMEM/ 5 mM glucose for 4 h in order to mimic conditions closer to the metabolic situation in vivo. To examine the glycogen content under conditions of reduced oxygen concentration (hypoxia stress), cells were incubated in an oxygen-regulated incubator with 1% O_2_, 5 % CO_2_ and 94 % N_2_ at 37 ° C for 4 h. Control cultures were kept under normoxic conditions (21 % O_2_). To study the influence of oxidative stress, hydrogen peroxide was added to the medium to a concentration of 500 μM and the cells kept for 30, 60 or 120 min, respectively, under these conditions. To examine the influence of lower pH values on the glycogen content (acidic stress), cultures were switched to 20 mM HEPES/Earle’s salts/ 5 mM glucose pH 7,4 (control) or 7,2 and 6,5 and incubated on a 37 °C water bath for 30 min.

After the experiments, the medium was removed, the plates quickly frozen at – 80 °and stored until glycogen determination.

### Cell viability

Trypan blue exclusion test was applied to determine cell viability.

### Animal studies

All experiments involving animals were carried out according to the law of animal experimentation issued by the German parliament (“Tierschutzgesetz”) and to the European Communities Council Directive.

### Statistics

Data are expressed as individual values and include determinations made in one experiment with n = 3 to 6. At least three replicates were made for one experiment. p values were obtained by student’s unpaired two-tailed t-test using the GraphPad Prism software.

## Acknowledgement

The authors thank Joan J. Guinovart for the kind gift of a test sample of glycogen synthase antibody.

## Abbreviations

*The abbreviations used are*

GPBB: glycogen phosphorylase brain isoform;
GPMM: glycogen phosphorylase muscle isoform;
GPLL: glycogen phosphorylase liver isoform;
CNS: central nervous system;
PNS: peripheral nervous system;
DRG: dorsal root ganglia;
DMEM: Dulbecco’s modified Eagle’s medium;
FCS: fetal calf serum

